# Generating Multi-state Conformations of P-type ATPases with a Conditional Diffusion Model

**DOI:** 10.1101/2024.08.07.607107

**Authors:** Jingtian Xu, Yong Wang

## Abstract

Understanding and predicting the diverse conformational states of membrane proteins is essential for elucidating their biological functions. Despite advancements in computational methods, accurately capturing these complex structural changes remains a significant challenge. Here we introduce a computational approach to generate diverse and biologically relevant conformations of membrane proteins using a conditional diffusion model. Our approach integrates forward and backward diffusion processes, incorporating state classifiers and additional conditioners to control the generation gradient of conformational states. We specifically targeted the P-type ATPases, a critical family of membrane transporters, and constructed a comprehensive dataset through a combination of experimental structures and molecular dynamics simulations. Our model, incorporating a graph neural network with specialized membrane constraints, demonstrates exceptional accuracy in generating a wide range of P-type ATPase conformations associated with different functional states. This approach represents a meaningful step forward in the computational generation of membrane protein conformations using AI and holds promise for studying the dynamics of other membrane proteins.

**TOC Graphic:** 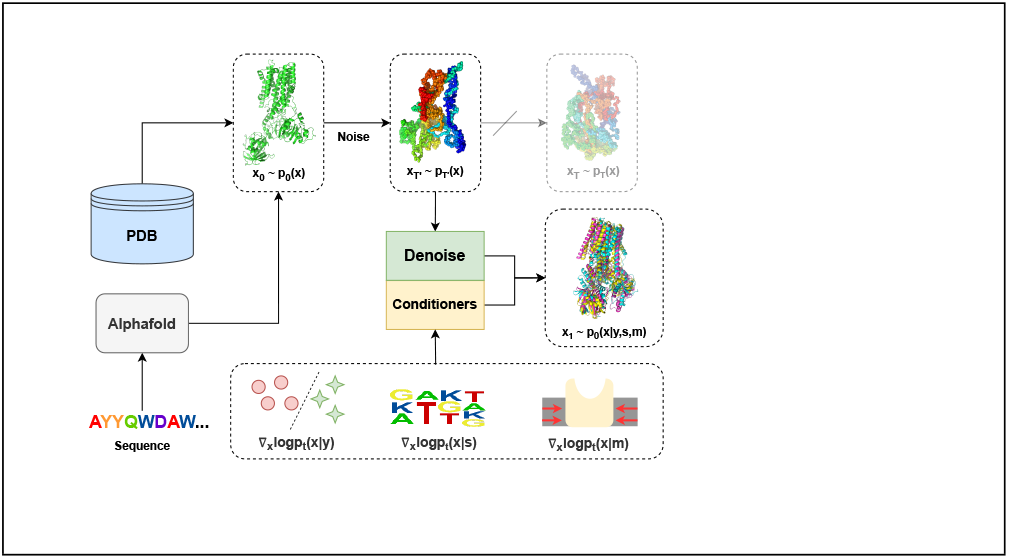

## Introduction

Proteins are inherently dynamic rather than static entities, with their functions deeply rooted in their capacity to change their conformations. These conformational changes are crucial as they allow proteins to interact with diverse ligands, substrates, and other macromolecules, thereby driving intricate biochemical processes. Thus the research on proteins in different states plays a vital role in elucidating their functional mechanisms, regulatory pathways, and interactions within biological systems. For instance, conformational flexibility is essential for enzyme catalysis, where substrate binding induces a shift in the enzyme’s structure, enhancing its catalytic efficiency.^1^ Similarly, protein-protein interactions often rely on the dynamic nature of protein conformations, enabling the formation of transient complexes necessary for signal transduction and cellular regulation.^2^ Analyzing these multi-state conformations is thus integral to comprehending the roles proteins play in biological systems.^3,4^

Currently, multiple methods exist for studying protein multi-state conformations, each with unique advantages and limitations. Experimental techniques such as X-ray crystallography, nuclear magnetic resonance spectroscopy (NMR), and especially cryo-electron microscopy (cryo-EM) which was bolstered by the recent “resolution revolution”,^5^ provide high-resolution structural snapshots but often demand substantial time, resources, and effort. On the other hand, *in silico* molecular dynamics (MD) simulations offer a state-of-the-art and feasible alternative, employing numerical solutions of Newtonian mechanics to explore protein conformational space.^6^ This approach effectively captures the microscopic processes of protein conformational changes, particularly in smaller systems, such as mini-protein folding and ligand binding.^7,8^ However, it struggles with larger protein systems with conformational changes occurring on timescales beyond tens of microseconds.^9^

To address these challenges, advanced enhanced sampling techniques have been developed, though they bring increased computational demands and complexity.^10–13^ Moreover, recent progress in deep learning has notably advanced structural biology, with initiatives to harness it for predicting multi-state protein conformations. ^14,15^ Methods such as finely tuned AlphaFold2 enable adjustments in structural templates, the depth of multiple sequence alignments (MSA), and the masking of MSA details to precisely characterize specific protein conformational states.^16–18^ For example, state-annotated templated GPCR databases assist AlphaFold2 in modeling both active and inactive states of GPCR at high accuracy.^19^ Nevertheless, despite these advancements, these approaches continue to grapple with issues related to interpretability or stability across diverse conditions.

Generative AI, exemplified by ChatGPT, has emerged as a focal point in recent research. By training neural networks to learn data distributions, this technology aims to generate new, realistic samples. While initially applied to computer vision, generative AI has recently made significant strides in life sciences, particularly within structural biology. As a prominent framework in generative AI, diffusion models are renowned for their capability to accurately represent complex distributions by emulating the underlying characteristics of the training data.^20–22^ In the context of biology, particularly in predicting protein structures, diffusion models offer a powerful approach due to their ability to model intricate conformational landscapes. For instance, the AlphaFold3 model replaces the structure model in AlphaFold2 with a diffusion model, enabling the prediction of joint structures of complexes, including proteins, nucleic acids, small molecules, ions, and modified residues, without excessive specialization.^23^ Chroma, another notable advancement, is a diffusion-based protein-generation model that enables the design of novel protein structures and sequences by integrating structured diffusion processes for protein backbones with scalable molecular neural networks.^24^ AlphaFlow and ESMFlow combine the flow matching framework with the AlphaFold2 and ESMFold models, accurately capturing conformational flexibility, positional distributions, and higher-order ensemble observables for proteins.^25^ Distributed graphormer (DiG) is a novel deep generative method aimed at approximating equilibrium distributions and efficiently sampling diverse and chemically plausible structures of molecular systems. ^26^ Diffold transforms AlphaFold2 into a diffusion model, achieving efficient sampling comparable to MD simulations.^27^ By introducing physical prior knowledge, ConfDiff incorporates a force-guided network with a mixture of data-based score models, enabling it to prioritize generating conformations with lower potential energy, which effectively enhances sampling quality.^28^ However, these models may still require further validation to ensure their predictions closely align with experimental data, and they have not been fine-tuned for membrane proteins. Membrane proteins present unique challenges due to their complex environments and interactions within the lipid bilayer, which are not always adequately captured by current models. Additionally, these generative models may struggle to generate structures in specific conformational states, which is crucial for understanding the functional mechanisms of proteins. This highlights important areas for future research and development in the application of generative AI in structural biology.

Building on the impressive achievements of generative AI in structural biology, we have developed a method utilizing the diffusion model to predict multi-state protein conformations. Inspired by the concept of heating and annealing, we propose a forward conditional-backward process for the diffusion model. By adjusting the duration of the noise scale, we can modulate the generated conformational space, thereby controlling the degree of conformational changes. Using the Bayesian theorem, we decompose the score function of the backward process into multiple conditioners, enabling precise control over the gradient direction and guiding structure generation. This method is highly modular, requiring minimal training of our conditioners to accurately predict multi-state conformations of membrane proteins, thus enhancing our ability to model complex biological systems.

In this study, P-type ATPases^29^ served as the model system to validate our methodology. We constructed a dataset, trained a classifier, and developed membrane constraint strategies specifically tailored for membrane proteins. These strategies provide essential contextual cues that enhance the prediction accuracy of membrane protein structures, thereby elevating the biological relevance of our computational models. Consequently, we were able to accurately generate distinct conformations of P-type ATPases in specific functional states. By comparing these generated structures to the latest cryo-EM structures of hSPCA1,^30^ we found a high degree of similarity between our model-generated conformations and their experimentally observed counterparts in similar states. Additionally, our analysis of the structural domains revealed that the generated P-type ATPases authentically mirror the characteristic features of their respective states. We anticipate the broad applicability of our method to generate diverse structures for a wide range of membrane proteins.

## Preliminary

### Diffusion model on protein structure

The diffusion model can be unified under the Stochastic Differential Equation (SDE) framework.^31^ As shown in the Fig. 1, X ∈ R^*N×*3^ represents the backbone structure of a protein, namely the Cartesian coordinates of all heavy atom including C*α*, C, N, O; *t* ∈ [0, *T* ] denotes the time step, and *w* is the standard Brownian motion regarded as disturbance term. The functions *f* (*x, t*) and *g*(*t*) are the drift coefficient and diffusion coefficient, respectively. The term ∇_*x*_log*p*_*t*_(*x*) is referred to as the score function, where *p*_*t*_(*x*) represents the distribution of the protein structure at time t. Protein structure can be sampled during inference utilizing either an empirical distribution^32^ or predicted through a neural network.

**Figure 1.**
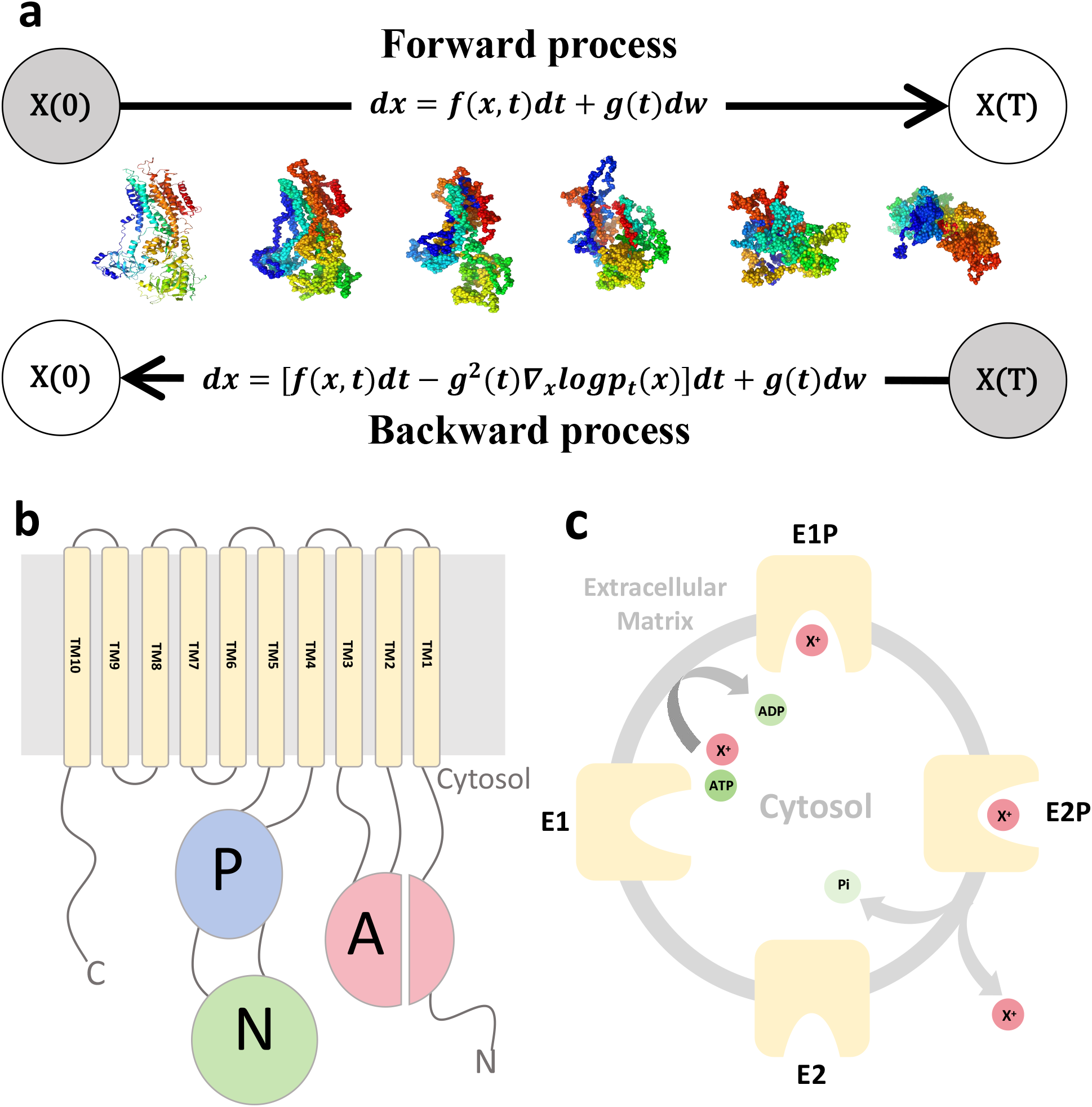
Diffusion model architecture and P-type ATPase conformational cycle. a) Illustration of diffusion model on protein structure generation in the form of a stochastic differential equation with noise space on the right and data space on the left. b) Schematic representation of the topology of the core subunit of a P-type ATPase. The actuator (A) domain is colored in red, the phosphorylation (P) domain in blue, and the nucleotide-binding (N) domain in green. (c) Illustrates the typical conformational cycle of a P-type ATPase with a cation *X*^+^ pump as an example, showing the sequential changes that occur during its conformational cycle.

The forward process transforms the native protein conformation *x*(0) into a noisy, disordered conformation *x*(*T*) by adding noise. For convenience, *p*_*T*_ (*x*) is chosen to be a simple, easy-to-sample distribution (usually Gaussian distribution). The backward process, derived from the given forward process, gradually denoises the noisy conformation *x*(*T*) back into the physically reasonable protein conformation *x*(0). The score function in this framework plays a key role in generation, which is often trained through deep learning. Utilizing the exponential-logarithm map, the process can be discretized similarly to the Euler–Maruyama step in Euclidean space as a geodesic random walk.^33^

To add conditions into the backward process, it is necessary to replace ∇_*x*_log*p*_*t*_(*x*) with ∇_*x*_log*p*_*t*_(*x*|*y*),^20^ where y represents the given condition. The precise ∇_*x*_log*p*_*t*_(*x*|*y*) can guide the gradient of structural generation during the backward process to obtain the protein structure under the specified condition.

### Conformation and substrate transport cycle of P-type ATPases

The P-type ATPases are a class of membrane proteins widely present in eukaryotes, which utilize ATP hydrolysis to translocate various substrates across membranes. ^29,34^ These ATPases harness ATP energy to transport substrates spanning from protons to phospholipids and are present in every cell type across all domains of life. Despite their overall low sequence conservation, they share a common structural fold and mechanism (Fig. 1b).^35^ The transmembrane (TM) region typically consists of 10 TM helices, which are crucial for substrate recognition and binding. The nucleotide-binding domain (N) functions as an intrinsic kinase binding ATP within a conserved amino acid motif. The phosphorylation domain (P) contains reactive aspartate residues subjected to transient phosphorylation. The actuator domain (A), serving as an internal phosphatase, houses conserved amino acid motifs that trigger the dephosphorylation of the phosphorylated intermediate. Classified into five sub-families (P1–P5), each specializing in distinct substrates, P-type ATPases generally follow the Post-Albers cycle.^36^ This cycle involves transitions between two primary functional states (E1 and E2) and their phosphorylated forms (E1P and E2P). During each cycle, inward and outward transport regulation occurs through the opening and closing of cytoplasmic and exoplasmic pathways, enabling access to TM binding sites with specific affinities and selectivities for transported substrates (Fig. 1c).

## Materials and Methods

### Structural dataset construction

We constructed our dataset using experimental structures from the IPR023298 superfamily of P-type ATPases, based on InterPro databank classifications.^37^ A total of 321 P-type ATPase structures were retrieved from the Protein Data Bank (PDB) released before May 2023. We streamlined this data by retaining only one major *α*-subunit from each X-ray or cryo-EM structure with duplicated chains and removing heteroatoms and ligands for later analysis.

Each structure was labeled with state tags (see Table 2), classifying P-type ATPases into four major states—E1, E1P, E2P, and E2—based on experimental conditions. Despite the presence of additional functional states (e.g., E1-ATP, E1P-ADP), we simplified the classification for consistency. For instance, E1P and E1P-ADP were grouped as E1P, and E1 and E1-ATP were categorized as E1. Structures that couldn’t be clearly assigned or did not fit the typical Post-Albers cycle states, such as auto-inhibited states,^38,39^ were excluded. Additionally, E2Pi was considered an intermediate state between E2P (with 80% similarity) and E2 (with 20% similarity).

To refine the dataset, we calculated TM-scores for all structure pairs and created a TM-score matrix (see Supporting Information). This matrix was used for hierarchical clustering to correct any incorrect state classifications.

### Data augmentation using molecular dynamics simulations

To enhance and balance the dataset distribution across different states, and to ensure the physical plausibility of the augmented data, we employed all-atom MD simulations. Given that the conformational changes of P-type ATPases between states typically occur over time scales longer than typical MD simulations,^40^ the nanosecond-scale simulations used here are intended only to sample conformational states in local equilibrium, rather than capturing large-scale state transitions. To accurately represent the membrane environment of P-type ATPases and streamline the model preparation process, we developed a template-based protocol to model the membrane protein within a lipid bilayer effectively. This approach, which avoids building each model from scratch, is outlined in the pipeline shown in Fig. 2. Initially, four P-type ATPases were selected as templates: KdpFABC in the E1-ATP state,^41^ SPCA1a in *Ca*^2+^-bound E1-ATP and E2 states,^42,43^ and yeast P4-ATPase, Dnf1, in the E2P state.^44^ These templates, with representative sizes ranging from 675 to 1158 amino acids, were modeled using the CHARMM-GUI web server^45^ and subsequently used to construct atomistic models of other P-type ATPases of varying sizes by following our template-based protocol (see Table 1). Dioleoylphosphatidylcholine (DOPC) was used for modeling the lipid bilayer.

**Table 1:** Four template models used for effective modeling P-type ATPases on membranes in subsequent MD simulations.

| PDB ID | P-type ATPase | Template size (a.a.) | Model size (a.a.) | PBC box |
| --- | --- | --- | --- | --- |
| 7ZRG | KdpFABC in E1-ATP state | 675 | (603,800) | 13nm x 13nm x 16nm |
| 7YAH | SPCA1a in $Ca^{2+}$-bound E1-ATP state | 877 | (800,900) | 13.5nm x 13.5nm x 17nm |
| 1IWO | SERCA1a in E2 state | 994 | (900,1085) | 15nm x 15nm x 18nm |
| 7WHV | yeast P4-ATPase, Dnf1 in E2P state | 1158 | (1085,1294) | 16nm x 16nm x 18nm |

**Table 2:**
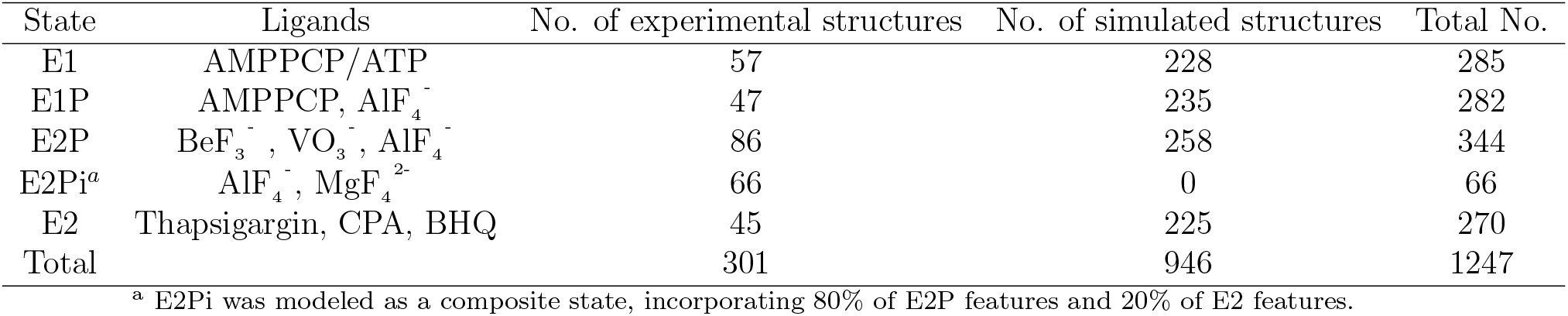
Summary of structure dataset of P-type ATPases.

**Figure 2.**
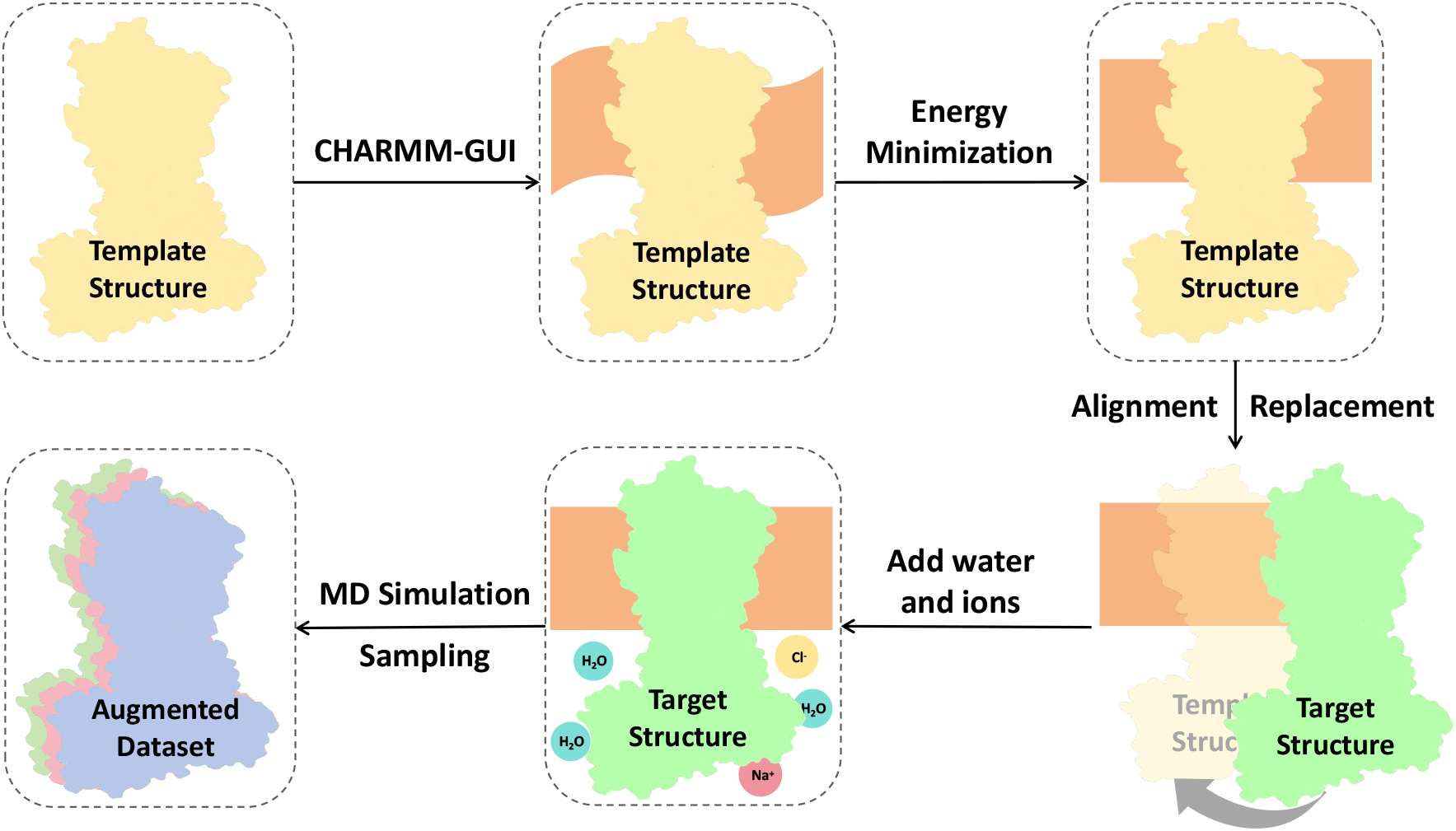
Workflow for atomistic modeling of P-type ATPases. This flowchart outlines the process of constructing atomistic models of various P-type ATPases. Template structures were used to generate models, which were then embedded in lipid bilayers and solvated. Energy minimization and all-atom molecular dynamics simulations were subsequently performed to refine and expand the structural dataset.

Next, energy minimization was performed using GROMACS (version 2022.5)^46^ to optimize the structures. For each structure in the dataset, PyMOL (version 2.5) was used to align and position it onto the membrane of the corresponding template, while DOPC molecules within 1.0 *° A* of the protein were removed to avoid steric clashes. Water molecules and 0.15M sodium and chloride ions were subsequently added to the model using the GROMACS gmx tool. After energy minimization, temperature equilibration, and pressure equilibration, MD simulations were conducted for 5-10 ns using the CHARMM36m force field. ^47^ The structures were uniformly collected along the MD trajectories at intervals of 1 ns per frame, with the number of MD structures for each state detailed in Table 2.

### Forward conditional-backward process

Inspired by the forward process and backward process of diffusion model, we propose a generation workflow for multi-state conformations of protein as shown in Fig. 3, which is based on a novel method named Forward Conditional-Backward Process. The process can be described by the following integral equation:

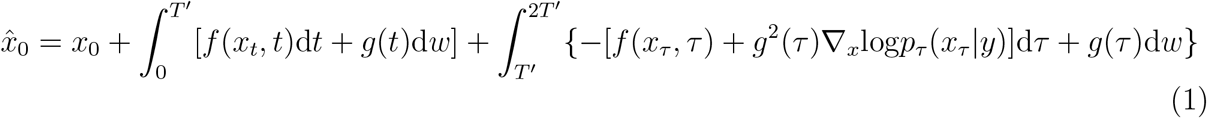

In Eq. 1,*τ* = 2*T*^*′*^ − *t*, y represents the specific conditions, 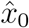 is the predicted conformation, *x*_0_ is the reference structure (obtained from experiment determination or structure prediction), and *T*^′^ ∈ (0, 1) denotes the noise duration. This equation describes the forward noise injection and backward denoising process toward the specific domain for a known structure. The noise time step *T*^′^ limits the scale of perturbation, thus preventing the complete elimination of information from the reference structure *x*_0_. Increasing *T*^′^ appropriately can enhance diversity but may require more reverse steps, while decreasing *T*^′^ can ensure the authenticity of the results but narrows the sampling space. Intuitively, it simulates the heating and annealing process. Here, the heating process aims to strengthen exploration, while the annealing process ensures authenticity.

**Figure 3.**
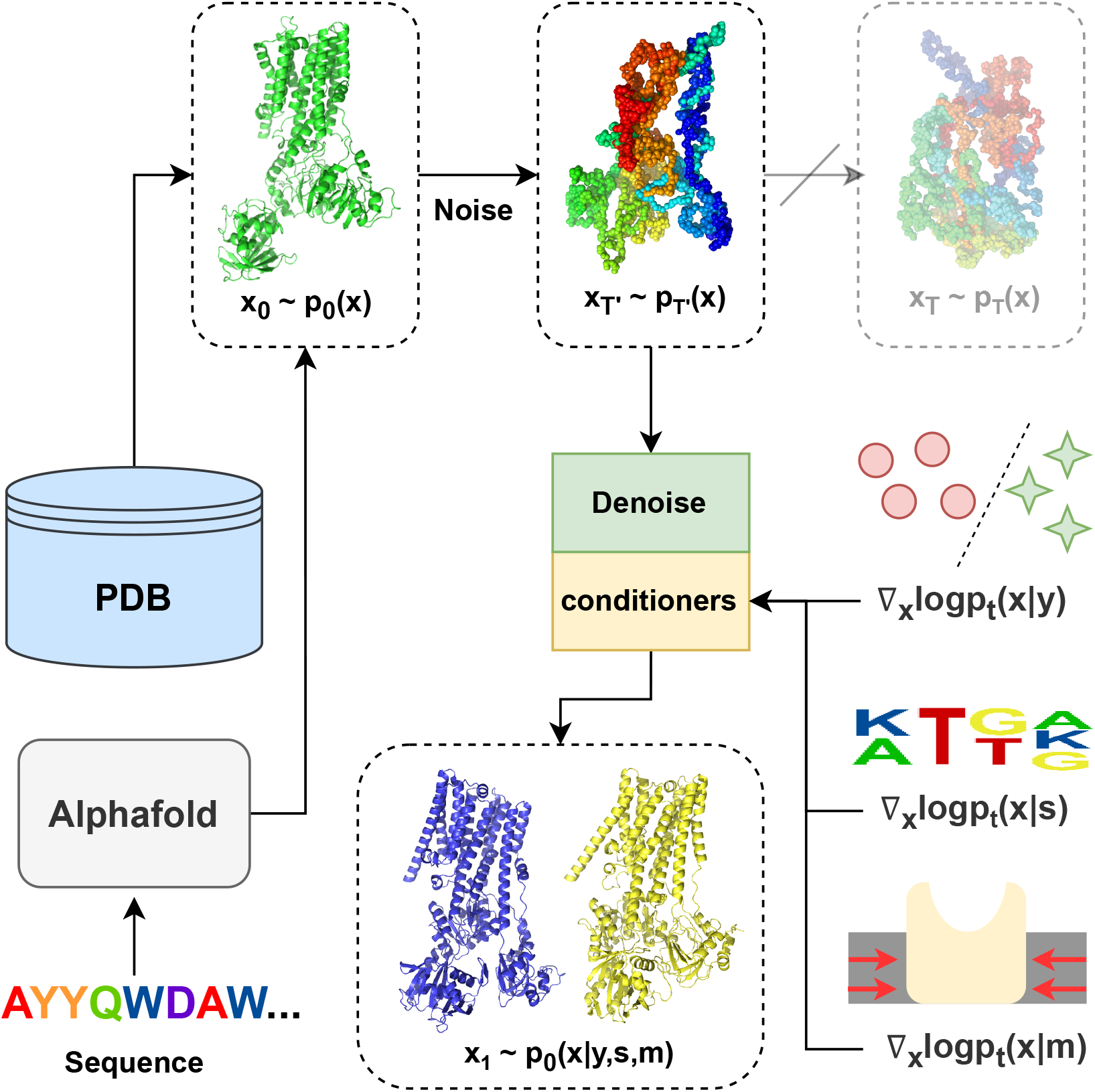
Workflow for multi-state conformation generation of Ptype-ATPases. Firstly, we need to obtain a reference structure observed through experimental methods or predicted by AlphaFold. This structure is then subjected to noise addition to x(t) during the forward process and subsequently denoised toward x(0) in a specific state through the conditional backward process with state classifier, sequence conditioner, and membrane constraint.

For the reverse process, we recommend the probability flow ODE scheme^31,48^ as follows:

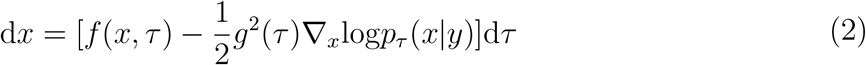

The ODE approach is a special case where the noise variance is zero. By eliminating the noise in the backward process, the model is reformulated as a continuous normalizing flow, which allows for efficient and precise likelihood computation using the adjoint method.^49^ This is reasonable when our goal is to improve model accuracy rather than diversity.

The score function in the formulation can treat arbitrary combinations of conditions if we model the joint event y as factorizing into independent sub-events *y*_1_, …, *y*_*N*_.^24^ Then we have the posterior score:

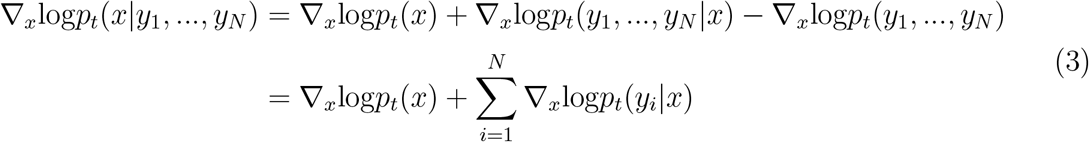

Thus the posterior score can be split into the sum of the origin score and different scores of conditioners. Specifically, the backward process involves three conditioners in our system: state conditioner, sequence conditioner, and membrane constraint. State conditioner refers to the specified state for generated conformations of protein. Sequence conditioner implies the constraint of the sequence given by the reference structure. The membrane constraint is specific to membrane proteins, representing the constraints imposed by the plasma membrane on the TM domain. The final posterior score should be:

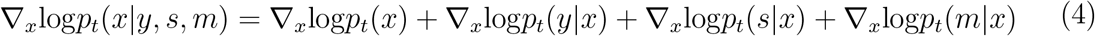

where x represents the structural coordinates, y represents the specified state, s represents the given sequence, and m represents the membrane constraint. The Eq. 4 includes the assumption that conditions y, s, and m are mutually independent. The final expression consists of four terms. The first term *p*_*t*_(*x*) and the third term *p*_*t*_(*s*|*x*) represent the stable distribution of the protein structure at time t and the distribution of the protein sequence given the structure x at time t, respectively. We utilize the models of these two terms from the open-source project Chroma to avoid unnecessary training. The second term, *p*_*t*_(*y*|*x*), represents the conformational state distribution of the protein structure x at time t. It is therefore referred to as the state classifier, which requires self-training. The final term, *p*_*t*_(*m*|*x*), represents the distribution of the protein structure x under membrane constraints at time t, which is thus referred to as the membrane constraint. The design of this term will be discussed later.

### Architecture and training of state classifier

The architecture is implemented based on PyTorch. Given the limited data, we selected a simple architecture comprising a graph neural network encoder (GNNEncoder), an attention pooling layer, and a multilayer perceptron (MLP), for classifier training.

In GNNEncoder, the diffusion time was first encoded using random Fourier featurization. Backbone coordinates were then utilized to extract 2-mer and chain-based distances and orientations.^24^ The sum of these components was passed to the later layer. A graph was constructed where heavy atoms on the backbone served as nodes, and interactions between atoms were treated as edges. Specifically, the construction of edges for each point was divided into two parts in the model. First, the 20 nearest points were identified to establish edges representing near-field interactions. Subsequently, 70 additional points were selected, with their selection probability weighted by the distance, to construct edges representing far-field interactions. The Graph Convolutional Networks architecture^50^ was adopted for message passing, which is defined as follows:

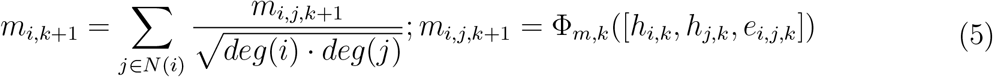

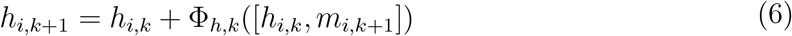

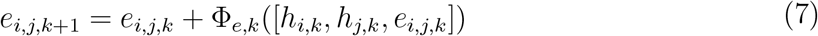

where *h*_*i,k*_ represents the SE(3)-invariant hidden representation of protein structure and nodes of the graph after k iterations; *e*_*i,j,k*_ represents the edge of the graph between *h*_*i,k*_ and *h*_*j,k*_ after k iterations; Φ_*m,k*_, Φ_*e,k*_ and Φ_*h,k*_ are various MLP at the k iteration; *N* (*i*) represents all the neighbors of node i; *deg*(*i*) represents the number of neighbors of node i. Our GNNEncoder model comprises four layers, with a node feature dimension of 512 and an edge feature dimension of 192. The Φ_*m,k*_ has two hidden layers, each with a hidden dimension of 128. The Φ_*h,k*_ includes two hidden layers, each with a hidden dimension of 256.

The Φ_*e,k*_ also has two hidden layers, each with a hidden dimension of 128. The activation function for all the MLPs was set to ReLU.

An attention pooling layer is required to aggregate feature representations based on importance weights and enhance model focus on relevant information for downstream tasks. The attention pooling is defined as follows:

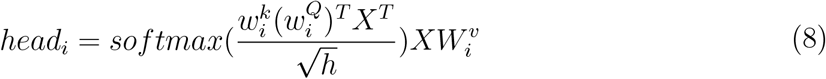

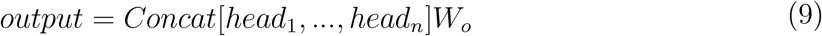

Here, m is the input dimension, h is the output dimension and n is the number of heads; 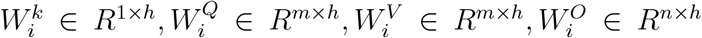 are the learnable matrices. In our model, we set n = 8 and h =512. Leveraging the attention mechanism helps the model to capture the relational information between residues.

Finally, the classification MLP has one hidden layer, with a hidden dimension of 64 and ELU as an activation function. The MLP outputs a four-dimensional vector representing four states for classification, with cross-entropy as the loss function to supervise the model learning.

We introduce the weights of the GNNEncoder from Chroma as our encoder section. Then we split the dataset into training (80%) sets and validation (20%) sets. The classifier predicts the following labels: E1, E1P, E2P, and E2. We used one-hot encoding for label processing and quantified the loss of each label prediction through cross-entropy. Subsequently, we optimized the parameters using the Adam optimizer^51^ with default momentum settings (beta = (0.9, 0.999)). We set the batch size to 8, linearly decreased the learning rate from 1e-5 to 1e-6, and trained for a total of 200 epochs. During training, we first uniformly sampled a time t ∈ (0,0.65) and sampled noise from the global covariance distribution.^24^ Then we injected this noise into the backbone coordinates and fed the results back to the classifier. Next, we predicted labels, calculated the loss, and updated the parameters.

### Membrane constraint

The lipid bilayer thickness of the cytoplasmic membrane is ∼4 nm,^52^ which can constrain membrane proteins through various interactions with the transmembrane domain, including hydrophobic interactions, electrostatic interactions, and hydrogen bonds.^53^ To simulate the interaction of membrane proteins with the plasma membrane in a realistic environment, it is necessary to abstract these constraints and apply them during the backward process of the diffusion model. Here, we propose two schemes: a reference-dependent constraint based on root mean square deviation (RMSD) and a reference-free constraint based on radius of gyration (Rg).

RMSD is a commonly used metric in structural biology, primarily for comparing the similarity between two different protein structures. In the RMSD scheme, we adopt a form of reconstruction guidance based on the optimal RMSD of the TM domain in the current denoised structure. Specifically, at time t, we define the membrane constraint *logp*_*t*_(*m*|*x*) as:

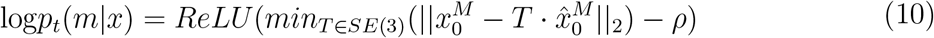

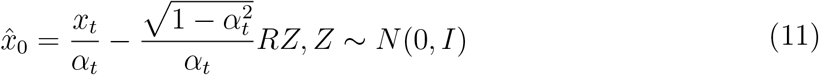

In Eq. 10 and Eq. 11,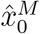 and 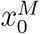 represent the TM domain of predicted structure at t=0 and the reference structure respectively. We set the z-axis as the normal to the plasma membrane, and define z_max_ as the maximum z-component of all atoms in the structure. Atoms within the range z ∈ [ z_max_-45, z_max_-5] are selected as 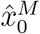 and 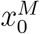 T is the optimal transformation matrix that minimizes the difference 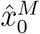 and 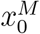. *ρ* is the threshold controlling the constraint range. This approach references the true structure to create a gradient that restricts the TM domain, effectively simulating membrane constraints. The advantage of this method lies in its simplicity and computational efficiency. Although this approach might incorporate information from the original state, the influence can be mitigated by adjusting weights.

Rg is a metric that describes the size of a protein in space, reflecting the average distribution of all atoms relative to its center of geometry. It is used to measure the compactness and spatial extension of the protein’s structure. In the Rg scheme, we constrain the diffusion of non-physical conformations in the TM domain by a penalty function based on the Rg of the TM region. Specifically, we define the constraint as *p*_*t*_(*Rg*^2^ *< k*|*X*_*t*_), where k is the threshold, and *Rg*^2^ is defined as follows:

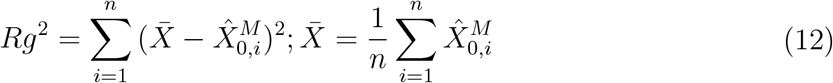

In Eq. 12, the 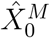 representing TM domain at time t=0 is predicted by *X*_*t*_. The advantage of this approach is that it avoids the bias towards specific state structures and only imposes constraints based on the threshold k. However, our calculations revealed that *Rg*^2^ follows a generalized *χ*^2^ distribution (see Supporting Information for details), indicating that the distribution of *Rg*^2^ does not have a closed-form solution. Consequently, specific values of *p*_*t*_(*Rg*^2^ *< k*|*X*_*t*_) can only be computed using simulation methods such as Monte Carlo simulations. With limited computational power, the results of Monte Carlo simulations are unsatisfactory, and gradient vanishing issues are likely to occur during backpropagation. Therefore, this approach is more suitable for detailed simulations when ample computational resources and time are available.

### Structure generation of P-type ATPases in specific states

In conditional generation models, adjusting the weight of the classifier is a common strategy.^54^ Modulating the classifier weight can balance output diversity and adherence to classification guidance. If the weight is too high, the generated structures may overly depend on the classifier’s guidance, resulting in homogeneous or unnatural content. Conversely, the generated content may deviate from the target category if the weight is too low. In practical applications, we do not strictly follow Eq. 4 but adopt the following form:

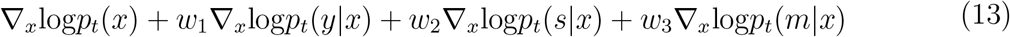

By adjusting various weights, we can control the diversity, reliability, and membrane constraint strength of the results. Ultimately, when generating protein conformations with the model, we used the following parameters: (1) Chroma parameters set step=500, *λ*_0_ = 10, *Φ* = 2, noise time step *T*^*′*^= 0.65. Additionally, to ensure quality over diversity, we applied the probability flow ODE in the backward process. (2) Classifier weight w1=1, gradient clipping value max norm=10. (3) Sequence conditioner weight w2=0.75. (4) Membrane constrainer using RMSD scheme, weight w3=0.1, threshold *ρ* set to 4. The generated structures currently contain only the backbone without side chains, which however can be modeled later using other algorithms, such as Modeller^55^ or GalaxyFill,^56^ if needed.

### Structure analysis and visualization

We used PyMOL for structural visualization; TMtools^57^ to calculate TM scores; Matplotlib to plot line graphs, violin plots, and bar charts; the R package Bio3D^58^ for PCA; and improved scripts from Bozzi et al.^59^ to calculate and average the distance difference matrices of all pairwise structures between two states.

## Results

### Structural dataset construction and its augmentation using molecular dynamics simulations

We constructed a structural dataset of P-type ATPases by curating PDB structures (released before 2023-05-01). Rigorous quality control was applied to ensure data integrity. To simplify the complex conformational space of P-type ATPases, we categorized structures into four primary states (E1, E1P, E2P, E2) and an intermediate state (E2Pi) which was defined to represent a hybrid of E2P and E2 states (see Methods). Structures lacking clear state assignment or deviating from the canonical Post-Albers cycle were excluded. The initial dataset, comprising 57 E1, 45 E2, 86 E2P, 47 E1P, and 66 E2Pi experimental structures, was augmented through explicit solvent all-atom molecular dynamics (MD) simulations using a protocol shown in Fig. 2, resulting in a final dataset of 1247 structures (Table 2).

### Development and validation of a functional state classifier

To initiate our analysis, we constructed a classifier capable of predicting protein states based on their structural information. As depicted in Fig. 4a, the model achieved a stable performance on the validation set, with a loss of 0.3 and an accuracy of 0.85, indicating its proficiency in discerning state-specific structural characteristics, even under noisy conditions. Nevertheless, the classifier demonstrated reduced precision in predicting the E1 state relative to other states. We attribute this disparity to the pronounced structural heterogeneity within the E1 state, encompassing diverse conformational states such as apo, calcium-bound E1, and E1-ATP. This structural variability likely hindered the model’s ability to identify common features characteristic of all E1 substates.

**Figure 4.**
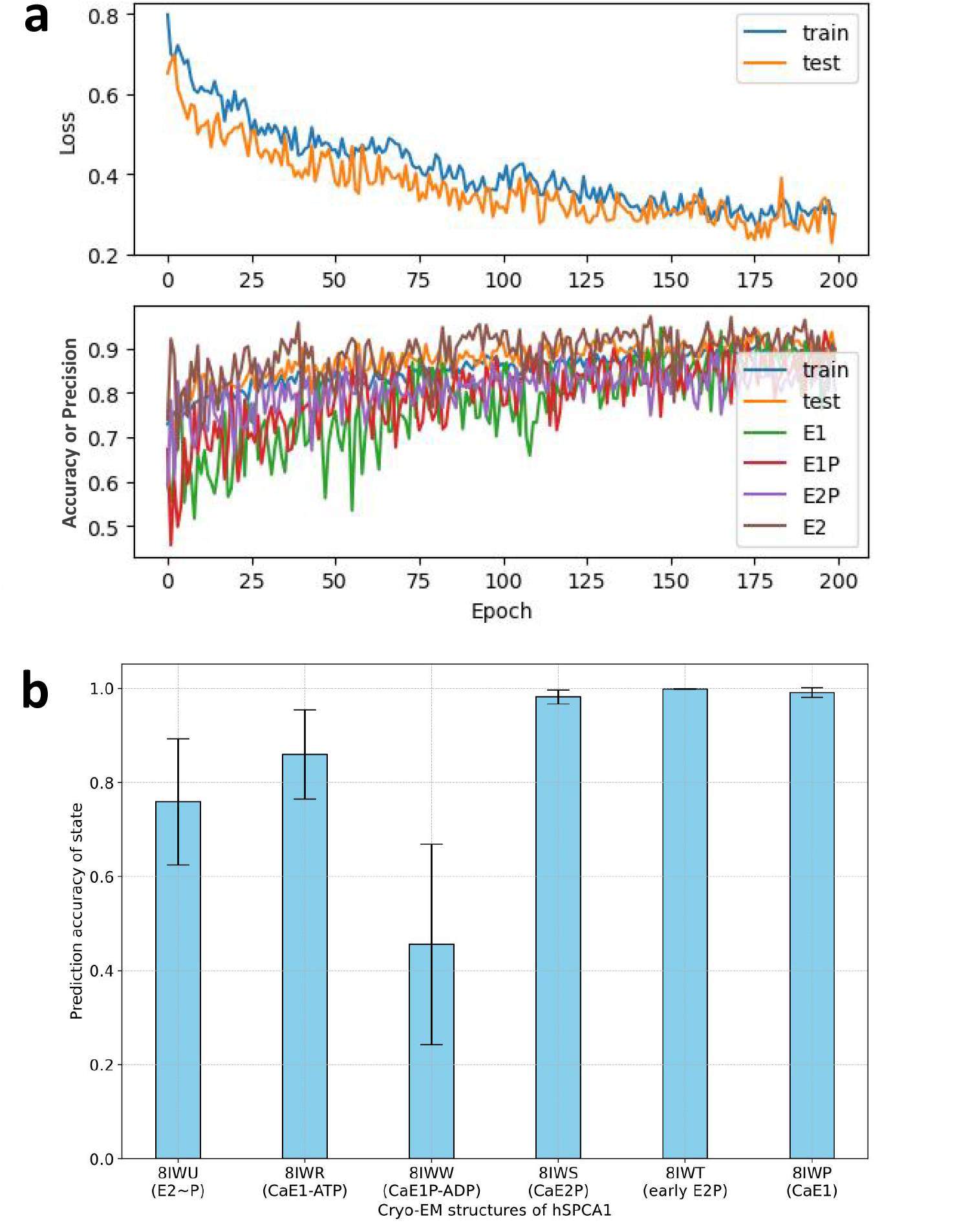
Training and evaluation of functional state classifier. (a) Learning curves illustrating the evolution of loss and accuracy during classifier training and evaluation on a separate test set. We calculated the accuracy for the training set and validation set and evaluated each conformational state with precision on the validation set. (b) Classifier accuracy on experimentally determined structures excluded from the training set. Error bars indicate the standard deviation in predictions.

To evaluate our classifier’s ability to generalize to unseen data, we applied it to a recently published dataset of human secretory-pathway calcium ATPases (hSPCA1) cryo-EM structures,^30^ which were excluded from the training process. hSPCA1 plays a critical role in maintaining *Ca*^2+^ homeostasis, and these cryo-EM structures represent a series of intermediate states, revealing a near-complete conformational cycle. We normalized the classifier’s output, a four-dimensional probability vector corresponding to the four functional states, using softmax. The highest probability was assigned as the predicted state. To mitigate the stochastic construction of graph during GNNEncoder, we repeated the prediction for each conformation 100 times.

As shown in Fig. 4b, the classifier accurately predicted the majority of conformations with accuracies ranging from 80% to 100%. However, the performance was significantly lower for the *Ca*^2+^-bound E1P-ADP state of hSPCA1 (PDB ID: 8IWW). We attribute this to the underrepresentation of early E1P states (including E1P-ADP) in the training data, which comprises only 2.1% of the E1P structure dataset. This finding underscores a potential bias in the classifier toward specific metastable states and highlights the need for more balanced training data to improve predictive accuracy for underrepresented states.

### Generation of conformations of hSPCA1 in specific states

To evaluate the capability of our model, we generated a series of conformations of hSPCA1 using our diffusion model, starting with the cryo-EM structure of hSPCA1 in the *Ca*^2+^-bound E1 state (CaE1, PDB ID: 8IWP) as the input structure. By altering the classifier’s targets, we directed the model to sample 15 times for each of the four states: E1, E1P, E2P, and E2, resulting in a total of 60 protein conformations. The structures were generated using a single NVIDIA GeForce RTX 3090, with each structure taking approximately 10 minutes to generate, amounting to a total of 10 hours. The representative structures are well-compacted (Fig. 5). Additional generation using the E2P state (PDB ID: 8IWS) as a reference structure is provided in the Supplementary Information (Figure S1).

**Figure 5.**
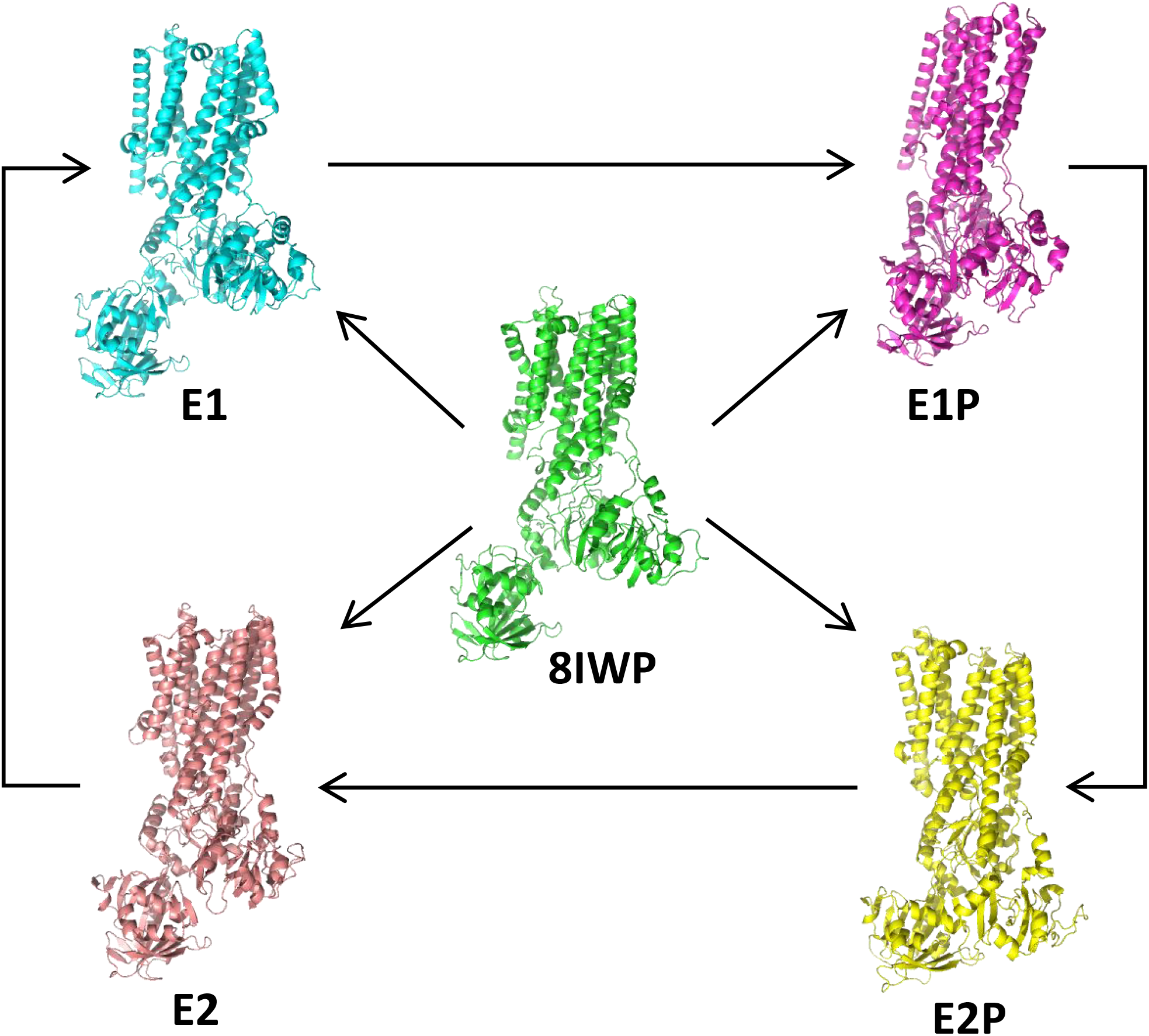
Diffusion Model-Generated Structures of hSPCA1. Our diffusion model utilized the cryo-EM structure of hSPCA1 in the *Ca*^2+^-bound E1 state (PDB ID: 8IWP) as a template to generate representative structures for four functional states. These states, E1 (cyan), E1P (magenta), E2P (yellow), and E2 (red), are depicted in cartoon representation.

To further assess the quality of these generated structures, we first used RMSD as a metric to examine the conformational similarity of the cytoplasmic domains between generated structures and the reference structure (CaE1, PDB ID: 8IWP), which exhibits high structural identity in different states. As shown in Fig. 6a, the RMSD for most domains remained around 2.0 *° A*, indicating stability during the generation process.

**Figure 6.**
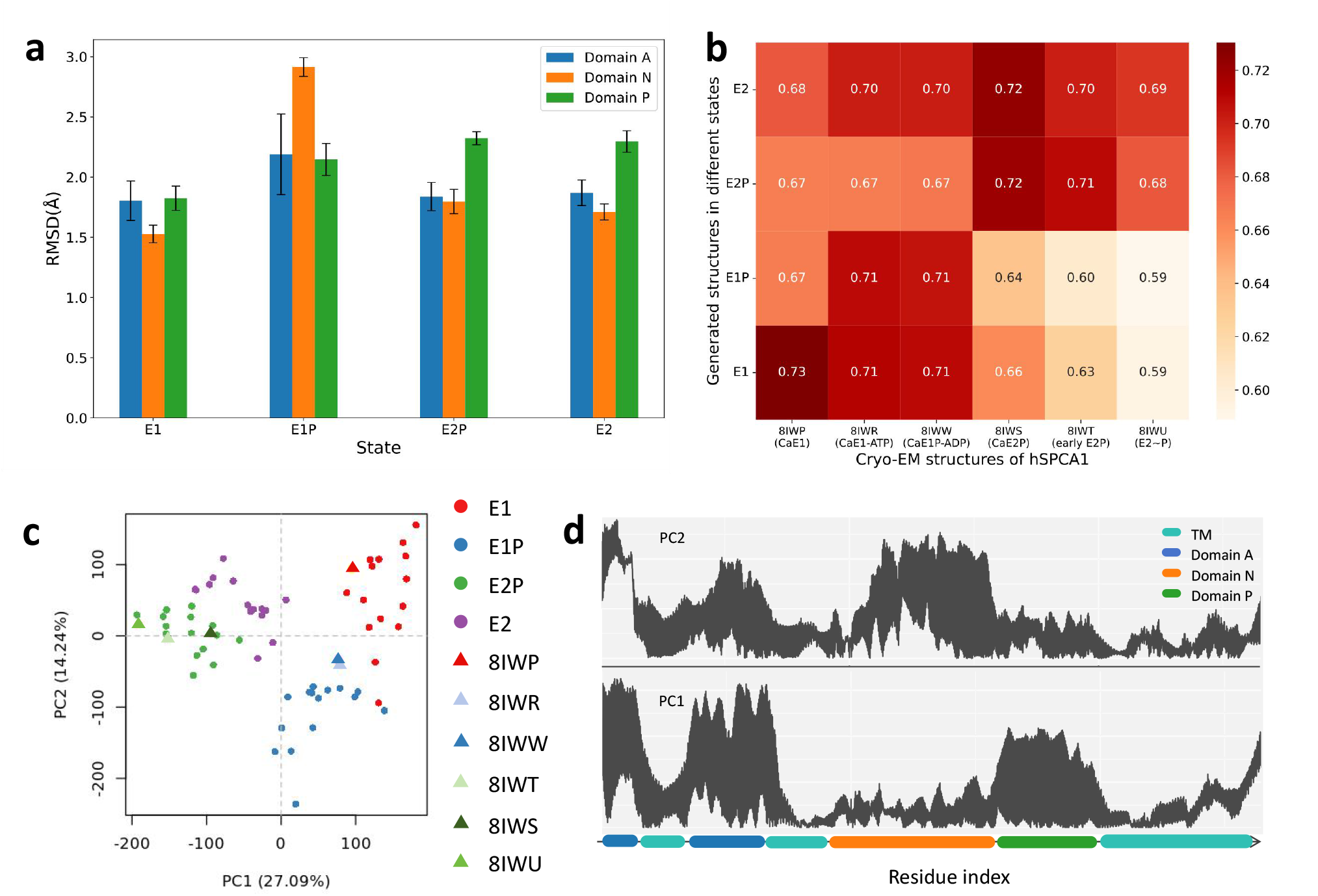
Structural diversity and conformational dynamics of generated hSPCA1 across different states. a) RMSD of individual domains (A: residues 136-248, N: 358-547, P: 548-687) between generated and reference (8IWP) structures. Bars are the mean and error bars indicate the standard deviation. b) TM score matrix comparing generated and experimental structures. Columns represent experimental structures, and rows represent averaged generated structures per state. c) Principal component analysis (PCA) of generated hSPCA1 conformations. States E1, E1P, E2P, and E2 are represented by red, blue, green, and purple dots, respectively. Experimental structures, including 8IWP (CaE1), 8IWR (CaE1-ATP), 8IWW (CaE1P-ADP), 8IWS (CaE2P), 8IWT (early E2P), 8IWU (E2∼P), are shown as triangles. Clustering of generated conformations with experimental structures is observed for E1, E1P, and E2P states, but not for E2 due to lack of experimental data. d) Line plot of PCA eigenvector, highlighting residue contributions to the principal component. The absolute values of the eigenvectors were used. The horizontal axis represents domains, with the vertical axis indicating weight magnitude.

We then compared the generated structures with the experimental structures of hSPCA1 which were considered the ground truth, using TM scores to evaluate precision and validity. The TM score matrix in Fig. 6b shows that the TM scores between the generated and experimental structures at corresponding states (8IWP, 8IWR, 8IWW, 8IWS, 8IWT, and 8IWU, representing the states of CaE1, CaE1-ATP, CaE1P-ADP, CaE2P, early E2P, and E2∼P, respectively) remained above 0.70. This indicates that our method reasonably predicts the structures in different states. Additionally, the lower TM scores between the generated structures and non-corresponding state structures reflect the state-specific characteristics of the generated structures. We also included an RMSD matrix in Supplementary Information Figure S2 to offer a comprehensive view of the structural differences and similarities.

To further explore structural differences and similarities, we performed PCA on the generated conformations. The resulting PCA plot in Fig. 6c shows distinct clustering of different states, indicating that the model effectively captured state-dependent conformational variations. We observed some diversity among the samples, particularly during the E1 state, characterized by domain activities in the cytoplasm, resulting in a broader sampling space. However, there is some overlap between different states, partly due to the inherent stochastic diffusion properties of the diffusion model. Despite using a probability flow model to eliminate noise during the backward process, the deep learning-based denoising network still exhibits some uncertainty. Additionally, heterogeneity among structures in the same state within the training set, such as E1, which can be further divided into metastable states like E1(apo) and E1-2Ca^2+^, results in some divergence in the classifier’s guidance of model generation. This can be attributed to the inherent stochasticity of diffusion models, training set heterogeneity, and the complexity of representing metastable states.

Subsequently, we used the top two eigenvectors of PCA, PC1, and PC2, to characterize the structural features (Fig. 6d). We observed that the weights exhibit a pattern closely related to the distribution of domains: the weights of the regions corresponding to the A, P, and N domains are generally at higher levels, indicating that these cytoplasmic domains are active during conformational changes and undergo significant dynamics. In contrast, most TM regions exhibit generally lower weights, suggesting that these TM helices are relatively stable during conformational changes. We also noticed that TM helices 1-2 and 9-10 have higher weights compared to other TM regions, suggesting they may be the main factors controlling P-type ATPase cavity changes, which is related to their vertical membrane displacement during the transition from E1P to E2P.^29^

### Generated structures capture the subtle conformational differences between functional states

Based on structural biology studies,^29, 60–62^ we selected specific features to depict structural details across various states, assessing whether the generated structures conform to reasonable conformational changes: (1) Distance between cytoplasmic domains, as significant displacements of these domains occur during phosphorylation and dephosphorylation phases; (2) Arrangement of TM regions, which is closely related to the binding and release of substrates.

The conformational changes of the cytosolic domains and the TM domain are shown in Fig. 7a and Fig. 7c. The distances between the centers of geometry of the cytoplasmic domains are shown in Fig. 7b. We observed that the distance between domains A and N is the longest in state E1, decreases during the E1 to E1P transition, and gradually increases in subsequent states to its original distance, forming a cycle. The small fluctuation in the distance between domains A and P during these conformational changes indicates a stable relative positioning of the A and P domains. The slight variations between domains P and N are likely due to peptide chain linkages connecting these domains. The distances in the reference structures further support the rationality of the generated structures.

**Figure 7.**
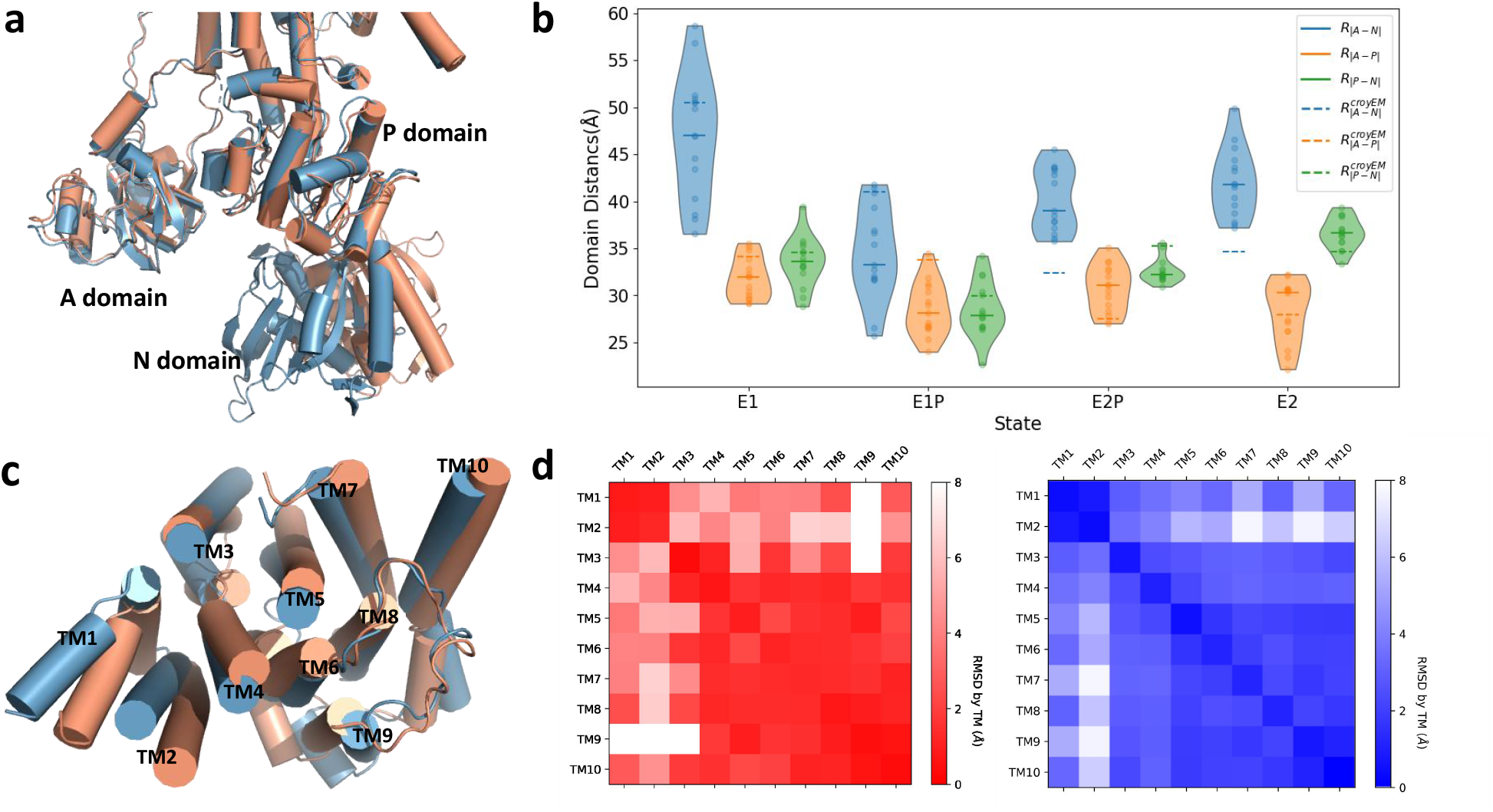
Conformational differences between the generated structures in functional states. a) Structural alignment of cytoplasmic domains. 8IWP (E1, light orange) and 8IWW (E1P, light blue) are aligned and compared, highlighting variations in the cytoplasmic A, P, and N domains. Alpha helices are shown as cylinders. b) Inter-domain distances within hSPCA1 cytoplasmic domains. Distances were calculated between domain centers of geometry, using 8IWP, 8IWW, 8IWS, and 8IWU as reference structures for E1, E1P, E2P, and E2 states, respectively. Caps at the top and bottom indicate the data range, with the solid line representing the median. c) Structural alignment of TM domains. 8IWS (E2P, light orange) and 8IWT (early E2P, light blue) are aligned, showing structural variations in the TM domain, including TM1-TM10. d) Distance difference matrices comparing TM helices of E2P and E2 state after aligning the entire TM domain. The left panel shows the real structure comparison using 8IWS (CaE2P) and 8IWT (early E2P) as references. The right panel depicts the average distance difference matrix calculated from generated E2P and E2 structures.

During the transition from E2P to E2, the TM regions rearrange to accommodate substrate loading. We used a distance difference matrix to analyze these rearrangements in the TM helices. As shown in Fig. 7d, the generated structures closely match the ground-truth structures. TM1-2, TM3-4, and TM5-10 display low internal RMSD, indicating stability within these helices, which may move as collective subdomains. In contrast, we observed substantial RMSD between TM1-2, TM3-4, and TM5-10, suggesting relative conformational movement between these helices, consistent with their role as the substrate loading site.^63^

### Ablation study of model components for optimized structure generation

To better understand the impact of key components (sequence conditioner, membrane constraint, and state classifier) on our model’s performance, we performed an ablation study by generating structures with different combinations of these components. These ablation results were evaluated from the perspectives of generation speed and quality to assess their effects on model performance. The time taken to generate the structures is used as an indicator of generation speed. We used the parameters mentioned in the subsection “Structure generation of P-type ATPases in specific states”. For the membrane constraint based on Rg, we employed Monte Carlo simulations with a sampling size of 100,000.

As shown in Table 3, the structural accuracy improves progressively when the sequence conditioner and membrane constraint are included, highlighting their critical role in stabilizing structure generation. Moreover, with the addition of the classifier, the TM score between the generated structures and the target structure in the E2P state (PDB ID: 8IWS) increases significantly, exceeding the TM score relative to the reference structure in the E1 state. This indicates that the classifier effectively guides the model in generating structures specific to target states.

**Table 3:** TM-scores and generation time for models with varying components in ablation study.

|  | Model1 | Model2 | Model3 | Model4 | Model5 | Model6 |
| --- | --- | --- | --- | --- | --- | --- |
| Sequence conditioner |  | ✓ |  | ✓ | ✓ | ✓ |
| RMSD-based membrane constraint |  |  | ✓ | ✓ | ✓ |  |
| Rg-based membrane constraint |  |  |  |  |  | ✓ |
| Classifier (E2P) |  |  |  |  | ✓ | ✓ |
| Time (s) | 83.84±1.45 | 188.16±1.50 | 142.54±2.70 | 256.13±2.46 | 481.40±18.32 | 698.50±26.65 |
| TM score using the CaE1 reference (8IWP) | 0.47±0.08 | 0.56±0.08 | 0.65±0.03 | 0.69±0.08 | 0.70±0.05 | 0.67±0.07 |
| TM score using the CaE2P reference (8IWS) | 0.50±0.09 | 0.54±0.08 | 0.64±0.03 | 0.66±0.07 | 0.72±0.05 | 0.71±0.06 |

Note: PDB 8IWP (CaE1) was used as the reference structure for models with different components, generating 15 structures for each model. The components include the sequence conditioner, membrane constraints based on RMSD or Rg, and a classifier labeled E2P. The results include TM-scores comparing the generated structures with PDB 8IWP (CaE1) and 8IWS (CaE2P), and the time required for generating individual structures, presented as the mean and standard deviation.

When comparing membrane constraints using RMSD and Rg, the Rg-based scheme does not provide an improvement in speed or structural quality. However, structures generated under the Rg-based constraint exhibit a lower TM score relative to the reference structure, indicating that this approach introduces less structural bias across different states.

### Comparison with template-based structure prediction using ColabFold

An alternative approach for generating conformations of known states is through homology modeling or structure prediction using a reference template. To compare our method with this approach, we tested the template-based prediction of hSPCA1 using ColabFold.^64^ For this comparison, we employed custom templates from *Ca*^2+^ ATPases, specifically PDB 5XA7 (E1), 1T5T (E1P), 7YAM (E2P), and 2C8L (E2). To limit the depth of the multiple sequence alignment (MSA) and enhance structural diversity, we set `max-extra-seq` to 64 and `max-seq` to 32. Additionally, three different random seeds were used for each prediction to introduce variability, selecting the top five ranked structures for each seed. This resulted in 15 structures per prediction and a total of 60 structures.

The results showed relatively low RMSD values for the A, N, and P domains in all predicted states (Fig. S5a), highlighting AlphaFold2’s accuracy in predicting individual domain structures. However, the generated E1 state exhibited nearly identical TM scores across different states, including 8IWP (CaE1), 8IWW (CaE1P-ADP), 8IWS (CaE2P), and 8IWT (E2P), suggesting limited ability to distinguish specific conformational states (Fig. S5b). Additionally, the generated E1P structures aligned more closely with CaE2P than the intended E1P. This is further corroborated by Fig. S5c, which uses PCA to quantify structural similarities among the predicted conformations. E2P and E2 states closely aligned with their experimental counterparts, while E1 and E1P exhibited greater variability.

Overall, our findings suggest that while template-based predictors like ColabFold/AlphaFold can generate high-confidence structures for individual domains, they struggle to accurately reproduce full structures with precise domain arrangements in specific functional states, such as E1 and E1P of hSPCA1, even when provided with appropriate structural templates. In contrast, our method, though slightly less precise in single domains, offers greater flexibility and specificity in generating multi-state conformations, making it a valuable tool for studying multi-domain proteins like P-type ATPases.

## Discussion and Conclusion

This study highlights the significant potential of diffusion models in predicting and generating diverse conformational states of membrane proteins, specifically focusing on P-type ATPases. Our approach, which integrates forward and backward diffusion processes with state classifiers and membrane-specific constraints, successfully captured the dynamic structural variability of these proteins. The ability to control the generation of conformational states through precise modulation of the conformational space is a notable strength, particularly when dealing with the complex and dynamic environments of membrane proteins. The high degree of similarity between our model-generated conformations and experimentally observed structures, such as those of hSPCA1, underscores the robustness and accuracy of this method.

The success of our approach in accurately reproducing the structural diversity associated with different functional states of P-type ATPases demonstrates its potential as a valuable tool in structural biology. By bridging the gap between static structural data from experimental techniques and the dynamic landscapes explored by MD simulations, our model offers a complementary approach to studying membrane protein dynamics.

However, there are challenges and areas for improvement that need to be addressed. The model’s reliance on existing datasets of experimental structures and MD simulations means that its performance is closely tied to the quality and diversity of the input data. Additionally, while our model has shown promise with P-type ATPases, further testing and refinement are required to generalize this approach to other membrane protein families, particularly those with more complex conformational landscapes.

An exciting avenue for future research is the possibility of combining this diffusion-based approach with traditional MD simulations and enhanced sampling techniques. By integrating the strengths of both methods, we could achieve a more comprehensive understanding of protein conformational dynamics. The diffusion model could be used to generate a wide range of initial conformations that are then refined and explored in greater detail using MD simulations. This hybrid approach could leverage the generative capabilities of diffusion models to sample diverse conformational states, while MD simulations could provide detailed insights into the temporal evolution and stability of these states. Such a combination could enhance the ability to explore conformational landscapes that are challenging to capture with either method alone, ultimately leading to a more detailed and accurate understanding of membrane protein function.

In conclusion, our work represents a meaningful step forward in the computational generation of membrane protein structures and provides a foundation for future advancements in the field. As generative AI continues to evolve, we anticipate that methods like ours will play an increasingly important role in unraveling the complexities of protein structures, ultimately contributing to developing new therapeutic strategies and advancing biological research.

## Supporting Information

- Detailed derivation and methodology of the radius of gyration scheme for membrane constraints.
- Analysis and figure of structural diversity and conformational dynamics of hSPCA1 generated using the reference structure of CaE2P (PDB ID: 8IWS) across different states.
- Summary table of the classifier’s hyperparameters.
- Figure for generation of conformations using ColabFold.
- Figure for Distance difference matrices comparing various states for both generated and experimental structures.
- Figure for structural alignment among different states for both the cytoplasmic and transmembrane domains.
- Figure for RMSD matrix (in unit of *Å*) comparing generated and experimental structures of hSPCA1 with reference structure 8IWP across different states.

## Author contributions statement

Y.W. conceived the experiment(s), J.T.X. conducted the experiment and analyzed the results. J.T.X. and Y.W. wrote and reviewed the manuscript.

## Data and Software Availability

The data, source code, related scripts, and parameter files used in MD simulation (.mdp file) are available at GitHub (https://github.com/36041255/PtypeATPaseGenerator).

## Acknowledgements

Y.W. acknowledges the financial support of the National Natural Science Foundation of China (No. 32371300), the Zhejiang Provincial National Science Foundation of China (No. LZ24C050003), and the National Key Research and Development Program of China (No. 2021YFF1200404). We thank our lab members, Keying Wang and Jiancheng Luo for their valuable insights and discussions.

## Supporting Information

### Membrane constraint scheme using radius of gyration

The objective function *p*_*t*_(*Rg*^2^ *< k*|*X*_*t*_) calculates the probability that under the given structure *X*_*t*_ at a specific time t, the radius of gyration of the final structure at time 0 is less than

k. The formula for calculating Rg is:

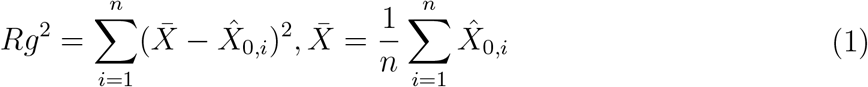

In this equation, *X*_0_ = [*X*_0,1_, *X*_0,2_, …, *X*_0,*n*_] represents the Cartesian coordinates of the heavy atoms in the structure at time t = 0. Here, the focus is on the positional relationships between the atoms rather than their physical and chemical properties. Therefore, for simplicity, all heavy atoms are considered to have the same mass. According to the forward process of the diffusion model:

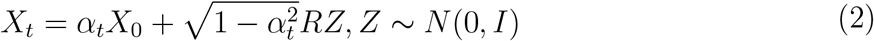

In this equation, *X*_0_ represents the true structure, and we can use this equation to estimate

*X* from 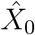 Rearranging the terms, we get 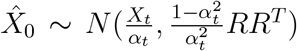 where R follows the globular covariance model:^**?**^

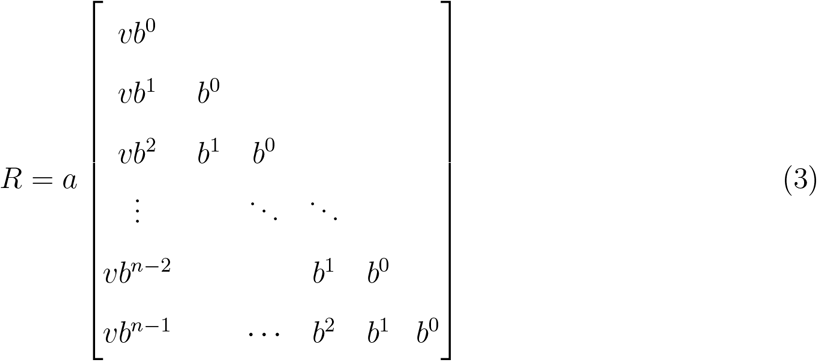

where n is the number of atoms, 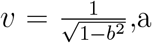 and b are constants greater than 0. Suppose *Z* = [*Z*_1_, *Z*_2_, …, *Z*_*n*_] are mutually independent standard normal distributions, then the noise *X*_*T*_ = *RZ*can be given by Z through the following formula:

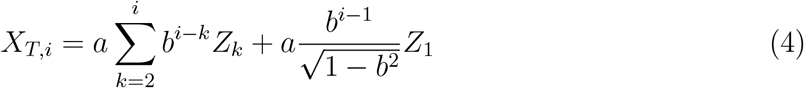

According to the property of normal distributions, it is known 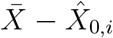 still follows a normal distribution. Now, let’s calculate the covariance between 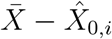 and 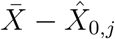 :

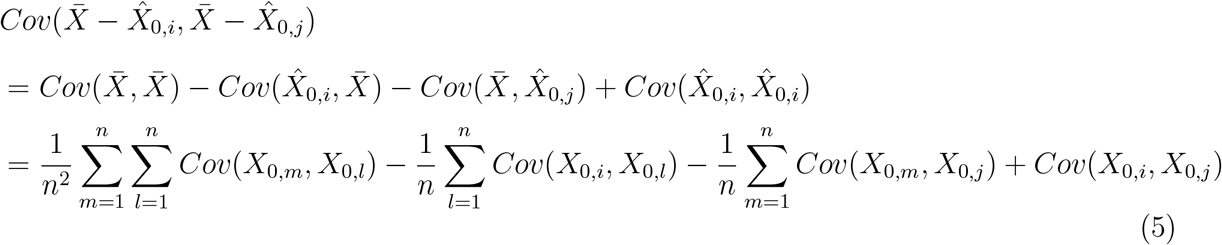

For any j ¡ i, we have:

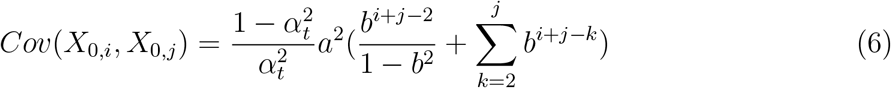

Obviously, 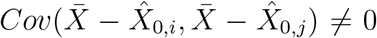 Therefore, *Rg*^2^ follows a generalized chi-squared distribution. The *p*_*t*_(*Rg*^2^ *< k*|*X*_*t*_) cannot be obtained in a closed-form solution and can only be approximated using methods such as Monte Carlo simulation.

**Table S1:** Classifier hyperparameters.

| Category | Hyperparameter | Value |
| --- | --- | --- |
| GNNEncoder | Number of the nearest neighbor edge | 20 |
|  | Number of the random edge | 70 |
|  | Nodes embedding dimension | 512 |
|  | Edges embedding dimension | 192 |
|  | Number of iteration for message passing | 4 |
|  | Message MLP dimension | 128 |
|  | Nodes MLP dimension | 256 |
|  | Edges MLP dimension | 128 |
| Attention pooling | Output dimension | 512 |
|  | Number of heads | 8 |
| MLP | Hidden dimension | 64 |
|  | Activation function | ELU |
| Training | Optimizer | Adam |
|  | Learning rate | linearly from 1e-5 to 1e-6 |
|  | batch size | 8 |
|  | Epoch | 200 |
|  | Noise scale | 0-0.65 |

## Generation of conformations of hSPCA1 with another reference structure in specific states

To evaluate the capability of our model, we generated a series of conformations of hSPCA1 using our diffusion model, starting with the cryo-EM structure of hSPCA1 in the *Ca*^2+^-bound E2P state (PDB ID: 8IWS) as the reference structure. By altering the classifier’s targets, we directed the model to sample 15 times for each of the four states: E1, E1P, E2P, and E2, resulting in a total of 60 protein conformations. The structures were generated using a single NVIDIA GeForce RTX 3090, with each structure taking approximately 10 minutes to generate, amounting to a total of 10 hours.

To further assess the quality of these generated structures, we first used RMSD as a metric to examine the conformational similarity of the cytoplasmic domains between generated structures and reference structure (8IWS), which exhibits high structural identity in different states. As shown in Figure S1a, the RMSD for most domains remained around 2 *Å*, indicating stability during the generation process.

We then compared the generated structures with the experimental structures of hSPCA1 which were considered the ground truth, using TM scores to evaluate precision and validity. The TM score matrix in Figure S1b shows that the TM scores between the generated and experimental structures at corresponding states remained above 0.72, except for the E1P state. This discrepancy may be due to 8IWR not being a representative E1P structure. As shown in Figure 4 of the main text, 8IWR was found to have a relatively lower accuracy. The result indicates that our method reasonably predicts the structures in different states. Additionally, the lower TM scores between the generated structures and non-corresponding state structures reflect the state-specific characteristics of the generated structures.

To further explore structural differences and similarities, we performed PCA on the generated conformations. The resulting PCA plot in Figure S1c shows distinct clustering of different states, indicating that the model effectively captured state-dependent conformational variations. However, We observed some diversity among the samples, particularly during the E1P state. Considering that this issue did not arise when 8IWP was used as the template structure, it is likely that this was caused by referencing the 8IWS structure in the E2P state.

Subsequently, we used the top two eigenvectors of PCA, PC1, and PC2, to characterize the structural features (Figure S1d). We observed that the weights exhibit a pattern closely related to the distribution of domains, which is similar to the generated structures with reference structure 8IWP: the weights of the regions corresponding to the A, P, and N domains are generally at higher levels, indicating that these cytoplasmic domains are active during conformational changes and undergo significant dynamics. In contrast, most TM regions exhibit generally lower weights, suggesting that these TM helices are relatively stable during conformational changes.

## Supplementary Figures

**Figure S1:**
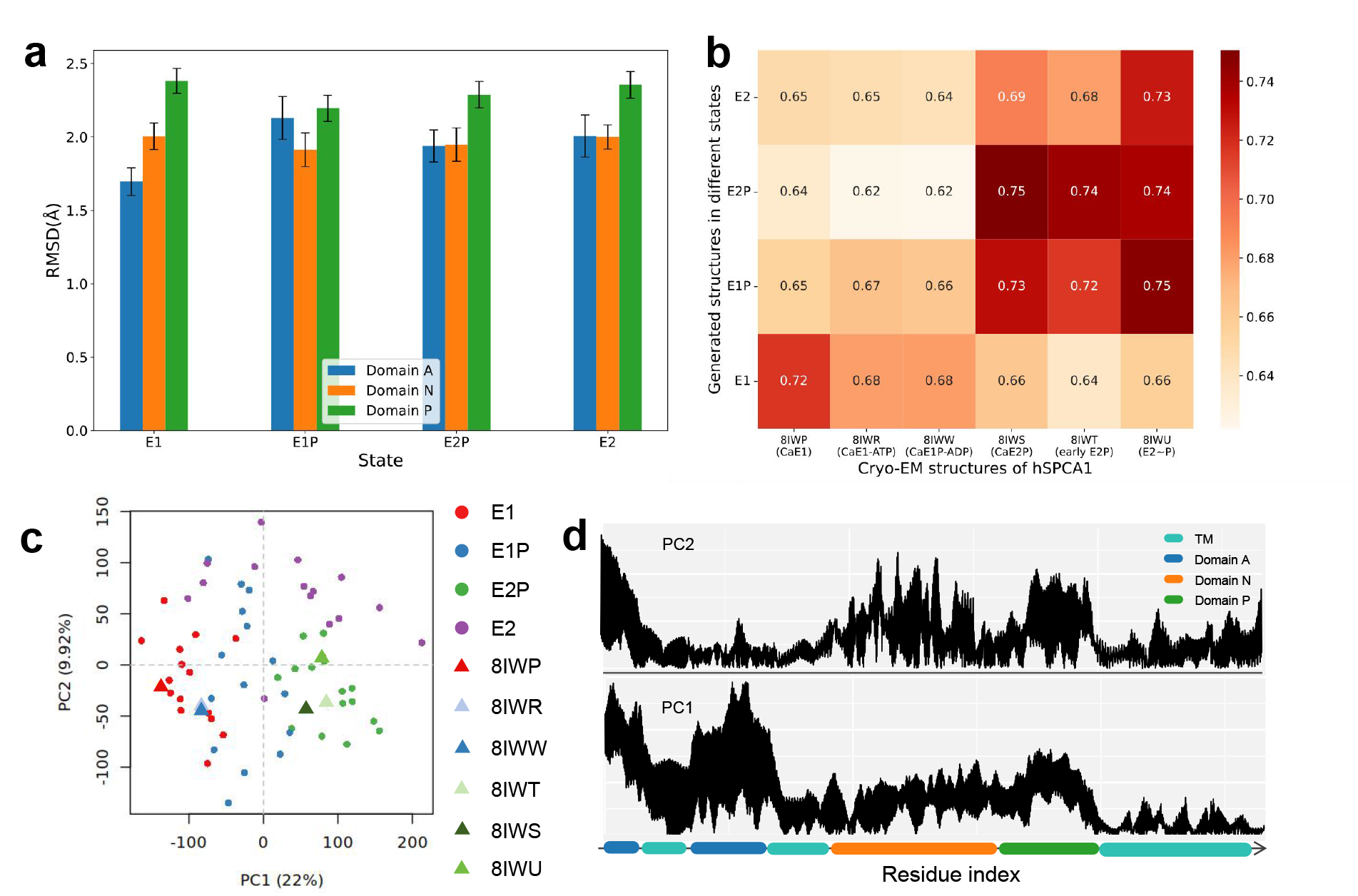
Structural diversity and conformational dynamics of hSPCA1 generated with reference structure 8IWS across different states. a) RMSD of individual domains (A: residues 136-248, N: 358-547, P: 548-687) between generated and reference (8IWS) structures. Bars are the mean and error bars indicate the standard deviation. b) TM score matrix comparing generated and experimental structures. Columns represent experimental structures, and rows represent averaged generated structures per state. c) Principal component analysis (PCA) of generated hSPCA1 conformations. States E1, E1P, E2P, and E2 are represented by red, blue, green, and purple dots, respectively. Experimental structures, including 8IWP (CaE1), 8IWR (CaE1-ATP), 8IWW (CaE1P-ADP), 8IWS (CaE2P), 8IWT (early E2P), 8IWU (E2∼P), are shown as triangles. Clustering of generated conformations with experimental structures is observed for E1, E1P, and E2P states, but not for E2 due to lack of experimental data. d) Line plot of PCA eigenvector, highlighting residue contributions to the principal component. The absolute values of the eigenvectors were used. The horizontal axis represents domains, with the vertical axis indicating weight magnitude.

**Figure S2:**
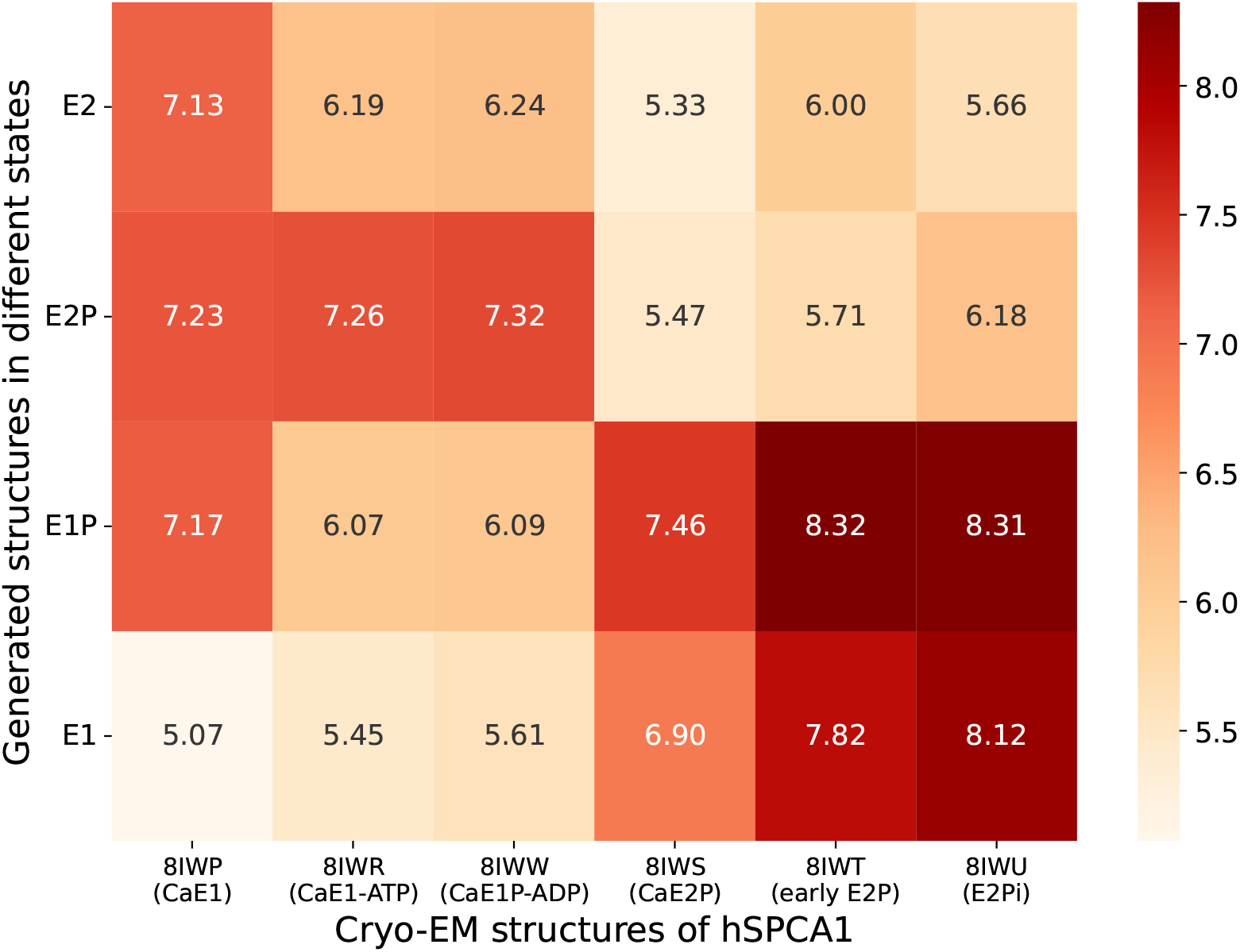
RMSD matrix (in unit of *Å*) comparing generated and experimental structures of hSPCA1 with reference structure 8IWP across different states.

**Figure S3:**
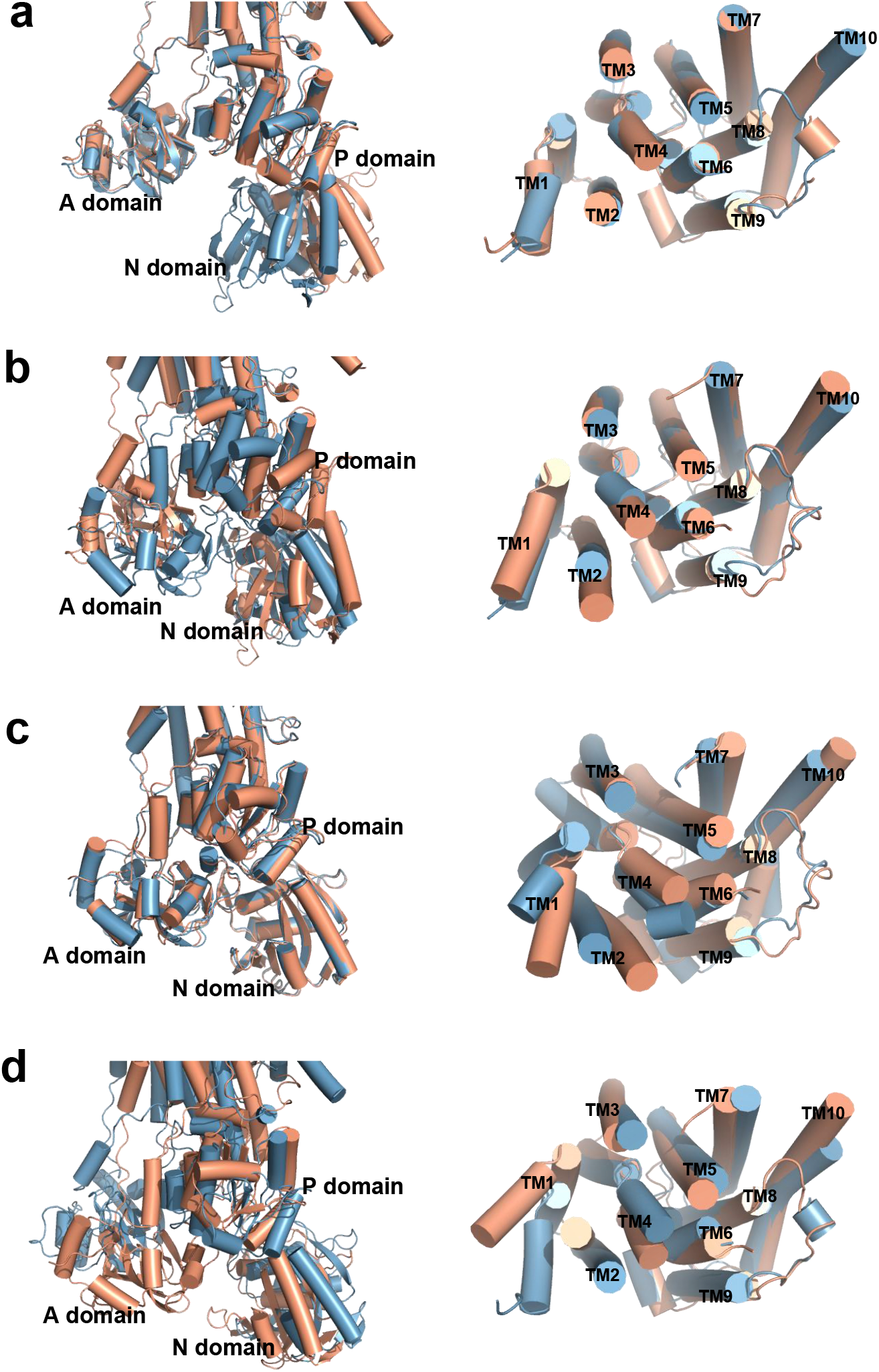
Structural alignment among different states of hSPCA1 for both the cytoplasmic and transmembrane domains. a) The alignment between 8IWP (CaE1) and 8IWW (CaE1P-ADP). 8IWP is represented in light orange, while 8IWW is in light blue. b) The alignment between 8IWW (CaE1P-ADP) and 8IWS (CaE2P), with 8IWW in light orange and 8IWS in light blue. c) The comparison between 8IWS (CaE2P) and 8IWT (early E2P), where 8IWS is light orange and 8IWT is light blue. d) The alignment between 8IWU (E2∼P) and 8IWP (CaE1), with 8IWU in light orange and 8IWP in light blue.

**Figure S4:**
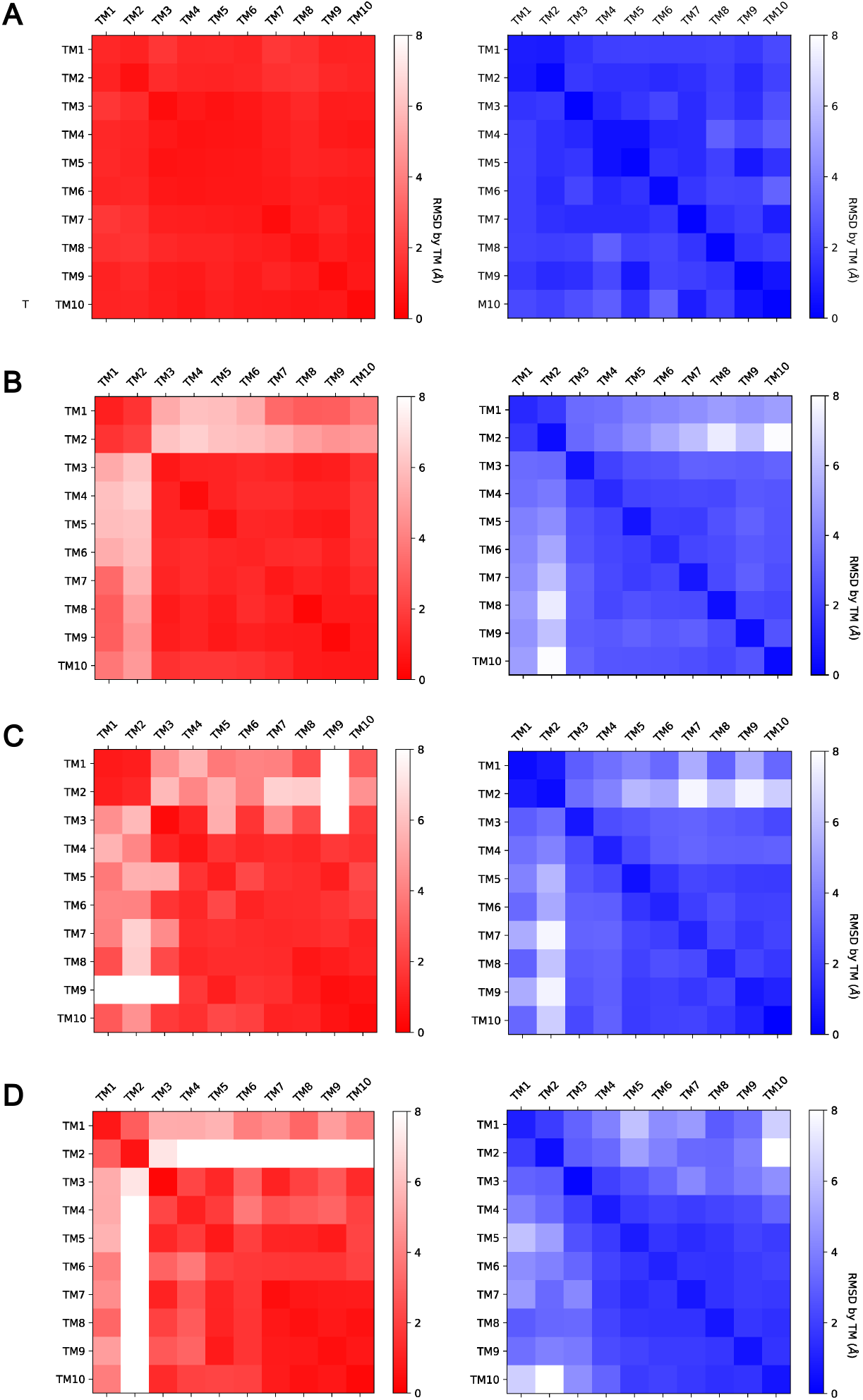
Distance difference matrix between different states. Structural differences between AI-generated states (blue matrices) and experimental structures (red matrices) of SPCA1 were analyzed. Each matrix represents the distance variations between corresponding transmembrane domains in the two structures. A-D): Each panel presents a comparison between a pair of AI-generated states (E1-E1P, E1P-E2P, E2P-E2, E2-E1) and their respective experimental reference structures (8IWP-8IWW, 8IWW-8IWS, 8IWS-8IWT, 8IWU-8IWP). The PDB codes in parentheses correspond to the experimental structures used as references. As the experimental structure of E2 is not yet available, the E2∼P structure (PDB 8IWU) was used as a reference for the E2-E1 comparison.

**Figure S5:**
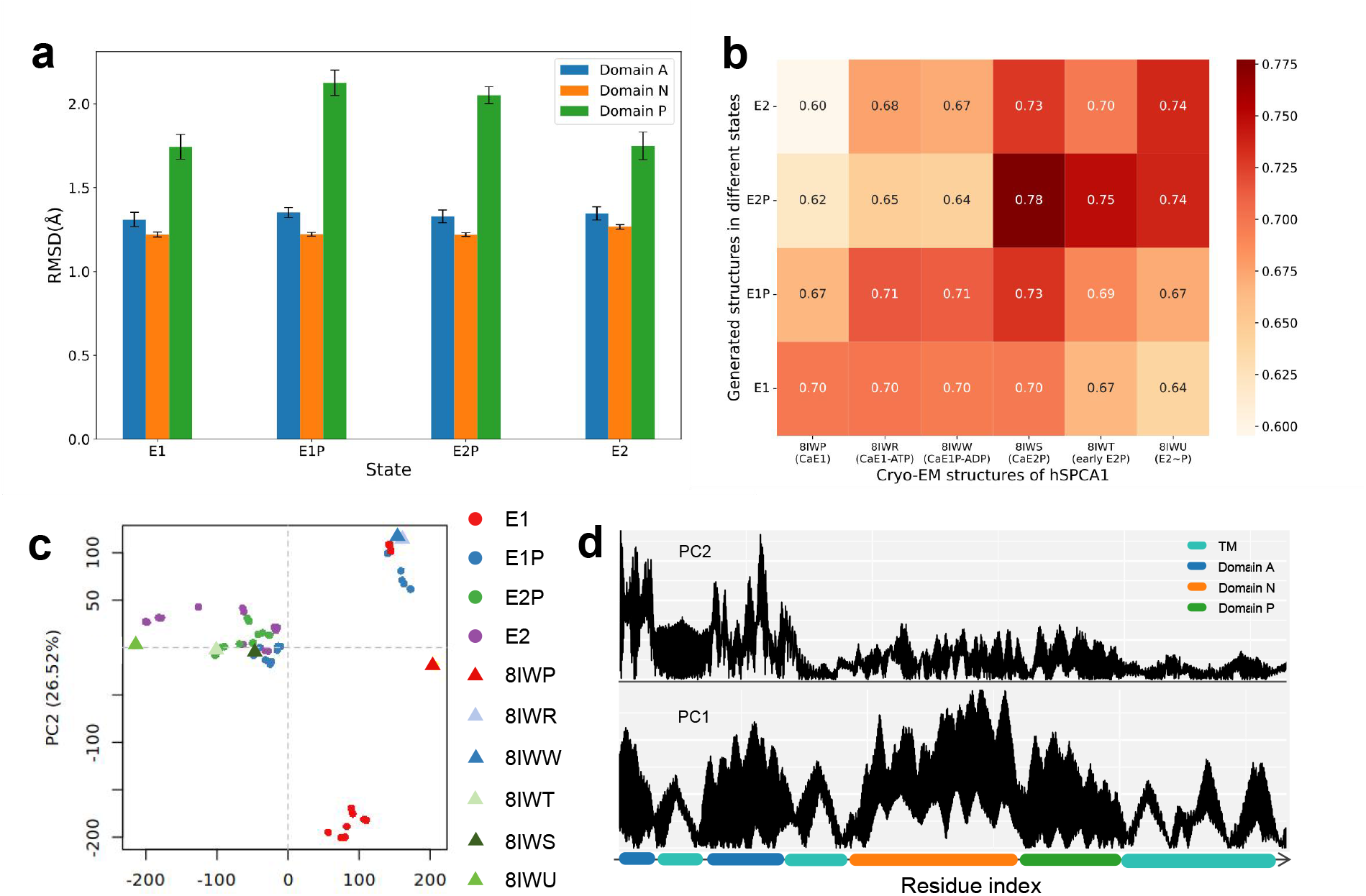
Structural diversity of hSPCA1 across different states predicted by template-based ColabFold. a) RMSD of individual domains (A: residues 136-248, N: 358-547, P: 548-687) between predicted and representative (8IWP, CaE1) hSCAP1. Bars are the mean and error bars indicate the standard deviation. b) TM score matrix comparing predicted and experimental structures. Columns represent experimental structures, and rows represent averaged predicted structures per state. c) Principal component analysis (PCA) of predicted hSPCA1 conformations. States E1, E1P, E2P, and E2 are represented by red, blue, green, and purple dots, respectively. Experimental structures, including 8IWP (CaE1), 8IWR (CaE1-ATP), 8IWW (CaE1P-ADP), 8IWS (CaE2P), 8IWT (early E2P), 8IWU (E2∼P), are shown as triangles. Clustering of predicted conformations with experimental structures is observed for E1, E1P, and E2P states, but not for E2 due to lack of experimental data. Some structures are too similar, resulting in some overlapping points. d) Line plot of PCA eigenvector, highlighting residue contributions to the principal component. The absolute values of the eigenvectors were used. The horizontal axis represents domains, with the vertical axis indicating weight magnitude.

